# A high-quality genome assembly and integrative data portal (*Phabase*) for the Mesoamerican black bean (*Phaseolus vulgaris* cv. Negro Jamapa)

**DOI:** 10.64898/2025.12.09.693262

**Authors:** Turgut Y. Akyol, Eber D. Villa-Rodriguez, Heladia Salgado, Emanuel Pacheco, Nancy Trujillo, Lavinia I. Fechete, Stig U. Andersen, Damien Formey, Jesús Montiel

## Abstract

Common bean (*Phaseolus vulgaris* L.) is one of the major legume crops worldwide, constituting a fundamental part of the diet in several countries. In central and south Mexico, the Mesoamerican Negro Jamapa variety is the one of most widely consumed cultivar, and it has notably served as a working model for functional genomics to address the molecular responses to stresses and during the nitrogen-fixing symbiosis with rhizobia. Despite this, the genome sequence of this landrace is not yet available, and the transcriptomic data generated by different research groups on this species are scattered. This prompted us to carry out a *de novo* sequencing, assembly and gene prediction for Negro Jamapa genome, using Hi-Fi PacBio technology and advanced genomic tools, respectively. Herein, we present a high-quality genome assembly for Negro Jamapa at chromosome level. The assembled genome is 522 Mb length, with an N50 of 45Mb, 89X coverage and BUSCO completeness of 98.4%, parameters that are considerably better than the available Mesoamerican reference genome. In parallel, we built an expression atlas of *P. vulgaris* from various tissues and growth conditions, by re-mapping publicly available RNA-seq and small RNA-seq data produced in recent years by distinct research groups. This analysis is available for the scientific community in the new user-friendly portal developed in this study, *Phabase* (https://phabase.ccg.unam.mx/). This portal includes several tools such as BLAST, gene and microRNA expression atlas, JBrowse, download section and integration with other *Phaseolus* spp. genomes.

**Significance statement:** In this study we report the first genome sequence of *Phaseolus vulgaris* Negro Jamapa, one of the most widely consumed bean varieties and a key experimental model for multiple research groups. We also developed Phabase (Phabase), an integrated database hosting this genome together with BLAST, JBrowse, and a comprehensive expression atlas. This resource enables functional genomics, comparative analyses, and accelerated breeding of common bean.

## Introduction

Legumes are essential sources of protein and micronutrients for human nutrition. Among all, the common bean (*Phaseolus vulgaris*) is the most important crop across Latin America, with two major domesticated gene pools: Mesoamerican and Andean (Bitocchi *et al*., 2012) (Schmutz *et al*., 2014). Common beans diverged into these two principal gene pools, approximately 0.1 to 0.2 million years ago (Bitocchi *et al*., 2013). The corresponding species were domesticated independently around 8,000 years ago (Schmutz *et al*., 2014) (Bitocchi *et al*., 2013) and show strong differences in their seeds, their adaptation range, and their resistance to abiotic and biotic stresses. The consequence of this dual domestication is an extensive genomic differentiation, such as lineage-specific alleles and structural variants, responsible for key agronomic traits (Bitocchi *et al*., 2013) (Gepts, 1998).

The Negro Jamapa, a Mesoamerican variety, was developed in 1957 by the Mexican Instituto Nacional de Investigaciones Forestales, Agrícolas y Pecuarias (INIFAP) through a structured plant breeding program, becoming a staple food across diverse regions of Mexico and serves as a principal source of protein and dietary fiber (Chavez-Mendoza and Sanchez, 2017). The resilience, adaptability, and combined nutritional and ecological qualities of Negro Jamapa have made it a key focus in agricultural research and breeding programs directed at improving yield stability, disease resistance, and nutritional quality (Espinosa-Alonso *et al*., 2006) (Palacio-Marquez *et al*., 2021) and also a major contributor to food security and cultural identity in the region.

The chromosome-scale reference genome for common bean was first generated for the Andean landrace G19833 (Schmutz *et al*., 2014). This was followed by the assembly of the BAT93 Mesoamerican cultivar genome (Vlasova *et al*., 2016), unfortunately still not at the chromosome level to date, with detailed transcriptomic profiles from multiple tissues and developmental stages, regrettably not easily accessible by the bioinformatic skill-lacking common bean community members. In addition to the common bean genotypes originating from Mesoamerican or Andean gene pools, whole-genome assembly of a European *P. vulgaris* accession was released (Carrere *et al*., 2023). More recently, to identify genomic regions that contribute to flowering responses in European cultivars, whole-genome resequencing was performed on 232 *P. vulgaris* accessions, mainly from Europe (Rendon-Anaya *et al*., 2025). Furthermore, the first telomere-to-telomere (T2T) genome assembly of the YP4 cultivar has been completed using PacBio High-Fidelity reads, ONT ultra-long sequencing, and Hi-C technologies, resulting in an assembly with exceptional completeness (BUSCO score: 99.5%) (Wang *et al*., 2025). Despite these advancements, existing genomic data remain fragmented or limited in annotation consistency. The genomic diversity of elite cultivars, particularly those widely used in breeding programs or as model genotypes in physiological studies, like Negro Jamapa, remains underrepresented.

Model legumes such as soybean, *Medicago truncatula*, and *Lotus japonicus* benefit from dedicated, integrative web portals, which unify genome assemblies, annotations, mutant resources, and expression data within a single platform (Mun *et al*., 2016) (Pecrix *et al*., 2018). Among others, these portals provide several tools such as genome browsers for visualizing genomic features at multiple scales, BLAST and sequence search tools for similarity analysis, synteny viewers for comparative genomics, gene expression atlases showing tissue-specific and condition-specific patterns. For common bean, there is still no equivalent, cultivar-aware environment that links a high-quality Mesoamerican genome with a comprehensive, re-analysed expression atlas and modern visualization tools. At the same time, the growing volume of transcriptomic, proteomic, and epigenomic data (Kim *et al*., 2015; Formey *et al*., 2015; Subramani *et al*., 2024) has created an urgent need for integrative bioinformatic platforms that allow comprehensive exploration of gene expression and regulatory networks in *P. vulgaris*. This gap is particularly limiting for Negro Jamapa, cultivar from which much of the functional genomics work and the RNA-seq data production for common bean has been performed. However, the absence of a tailored reference genome forces researchers to project data onto non-Mesoamerican assemblies, leading to less accurate and efficient sequence-based analysis experiments.

Previous efforts, such as gene expression atlases, provided valuable insight but were constrained by old sequencing technologies, lack of biological replicates, and outdated genome annotations (O’Rourke *et al*., 2014). A unified, user friendly, and dynamically updated resource is essential to fully exploit the wealth of omics information and to support data-driven hypothesis generation in this crop. To address this gap, here, we present the first high-quality genome assembly of the Negro Jamapa cultivar using long-read sequencing, complemented with a comprehensive genome annotation. Further, we developed an integrative online portal, *Phabase* which hosts the corresponding assembled genome, gene annotations, and associated expression data. The portal includes interactive tools such as a genome browser, BLAST search, and an expression atlas of genes and microRNAs. Our genome assembly is based on PacBio HiFi sequencing, which enables highly contiguous and accurate assemblies. Leveraging this technology for Negro Jamapa will offer the opportunity for precise research in the crop’s comparative and functional genomics, genomic features relevant to symbiosis, biotic and abiotic stress responses, and yield-related traits.

## Results

### Assembly and quality metrics

The raw data of the whole genome sequencing of common bean Negro Jamapa consisted of 4,487,410 HiFi circular consensus sequencing (ccs) DNA reads with an average length of 21,232 bp and 90.77% of the bases had a quality score greater than 30. The size of the assembled genome is 522,239,992 bp with a GC content of 36.31%, 89X mean coverage and 98.4% completeness according to BUSCO. In addition to the 11 chromosomes and the mitochondrial and chloroplast genomes **(Table 1)**. There were 113 scaffolds that were mapped to the unplaced scaffolds in the reference genome v2.0 with a total length of 17,728,932 bp. The assembly includes 1,294 unplaced scaffolds collectively spanning 113,563,414 bp.

**Table 1.**
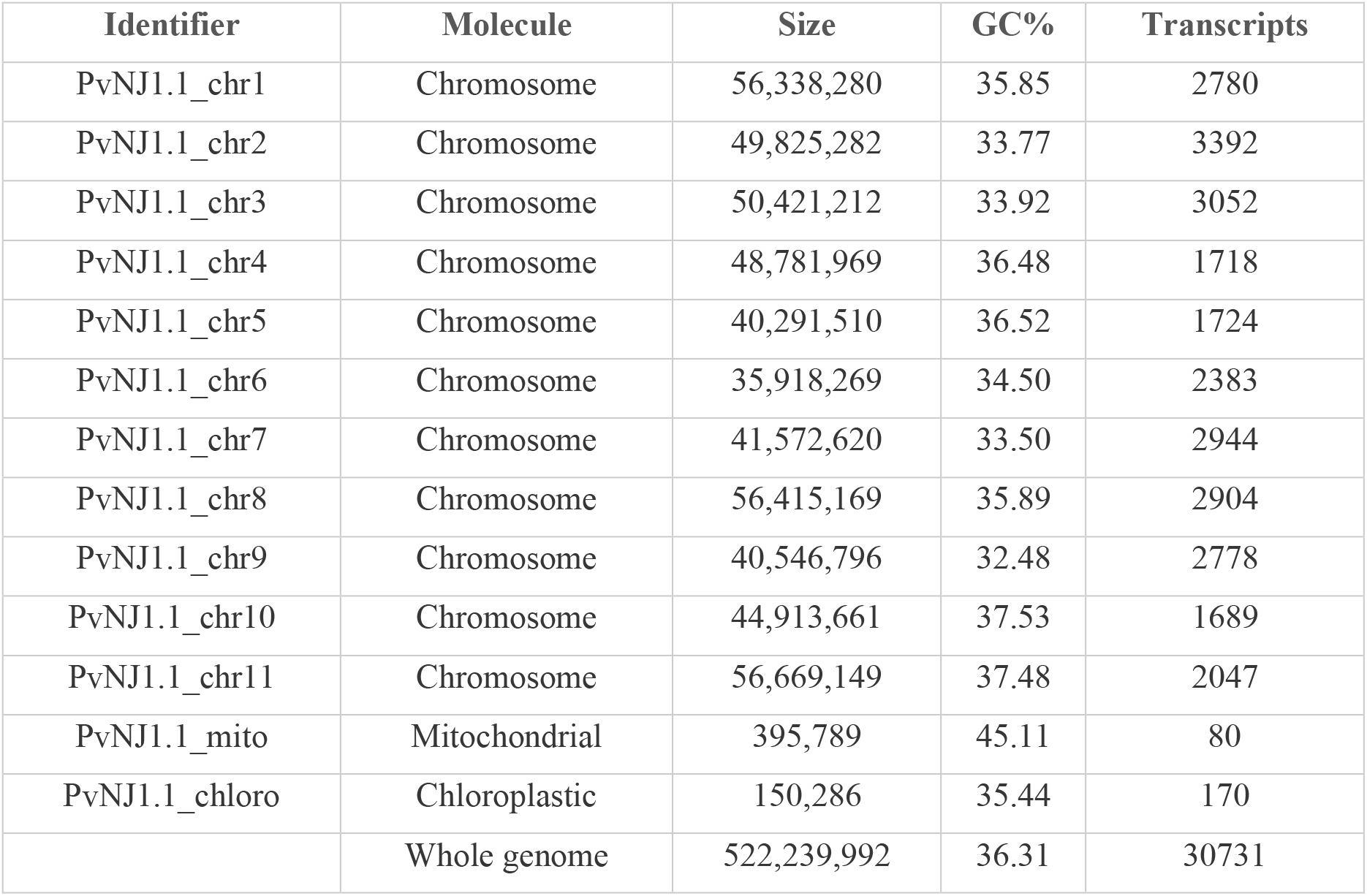
*P. vulgaris* cv. Negro Jamapa genome assembly and annotation statistics.

**Table 2.** Comparison of the *P. vulgaris* cultivars’ assemblies.

| Assembly | Genome size | Total ungapped length | Number of chromosomes | Number of scaffolds | Scaffold N50 | Genome coverage |
| --- | --- | --- | --- | --- | --- | --- |
| **Negro Jamapa** | 522.2 Mb | 522.2 Mb | 11 | 1420 | 44.9 Mb | 89x |
| **YP4** | 560.3 Mb | 560.3 Mb | 11 | 11 | 55.1 Mb | 56x |
| **INB_841** | 546.4 Mb | 546.4 Mb | 11 | 11,399 | 31.1 Mb | 43x |
| **G19833 v2.0** | 537.2 Mb | 531.5 Mb | 11 | 1,032 | 1.9 Mb | 83x |

### Comparative Genome Assembly and Annotation Metrics

The genome assembly of *P. vulgaris* cv. Negro Jamapa showed important improvement in contiguity and completeness compared with the reference genotype G19833 (Schmutz *et al*., 2014). The total size of the assembly was 522 Mb, which is close to the 531 Mb of G19833, but with a significantly higher N50 of 45 Mb instead of only 2 Mb in G19833. Sequencing coverage was also slightly higher in Negro Jamapa (89x) compared to G19833 (83x), supporting the high quality of the assembly. This improvement was mainly due to the use of long-read sequencing and better assembly strategies, allowing to reach almost chromosome-scale continuity.

We identified 27,635 protein-coding genes in the annotation of Negro Jamapa, similar to the 27,433 genes in G19833. BUSCO analysis showed completeness values of 98.4% for Negro Jamapa, slightly improved compared to the 97.7% estimated in G19833, confirming that both assemblies are almost complete and of excellent quality. In addition, the new annotation of Negro Jamapa contained more microRNA precursors annotated (270 vs. 226), which suggests a larger repertoire of small RNAs or a consequence of the higher number of publicly available sRNA libraries generated from Negro Jamapa samples. Together, these data indicate that the Negro Jamapa genome is an improved resource, with high contiguity, completeness, and annotation depth, useful for comparative and functional genomics studies in *P. vulgaris*.

### Genomic landscape and structural features

To better understand how the Negro Jamapa genome is organized, we analyzed the distribution of multiple genomic features across its eleven chromosomes **(Figure 1)**. The distribution of repetitive elements was uneven: certain chromosomes showed clear concentrations near their pericentromeric regions, consistent with the heterochromatic nature of these areas. The arrangement of annotated protein-coding genes revealed alternating regions rich and poor in genes, a pattern typical of the euchromatic and heterochromatic domains found in other legume genomes. When examining potential paralogous relationships and segmental duplications, we found that these events were particularly frequent among chromosomes 1, 8, and 11, hosts of more than 38% of the cumulated duplication length. This observation is consistent with our structural variation analysis, which identified several large translocations and highly diverged duplicated regions.

**Figure 1.**
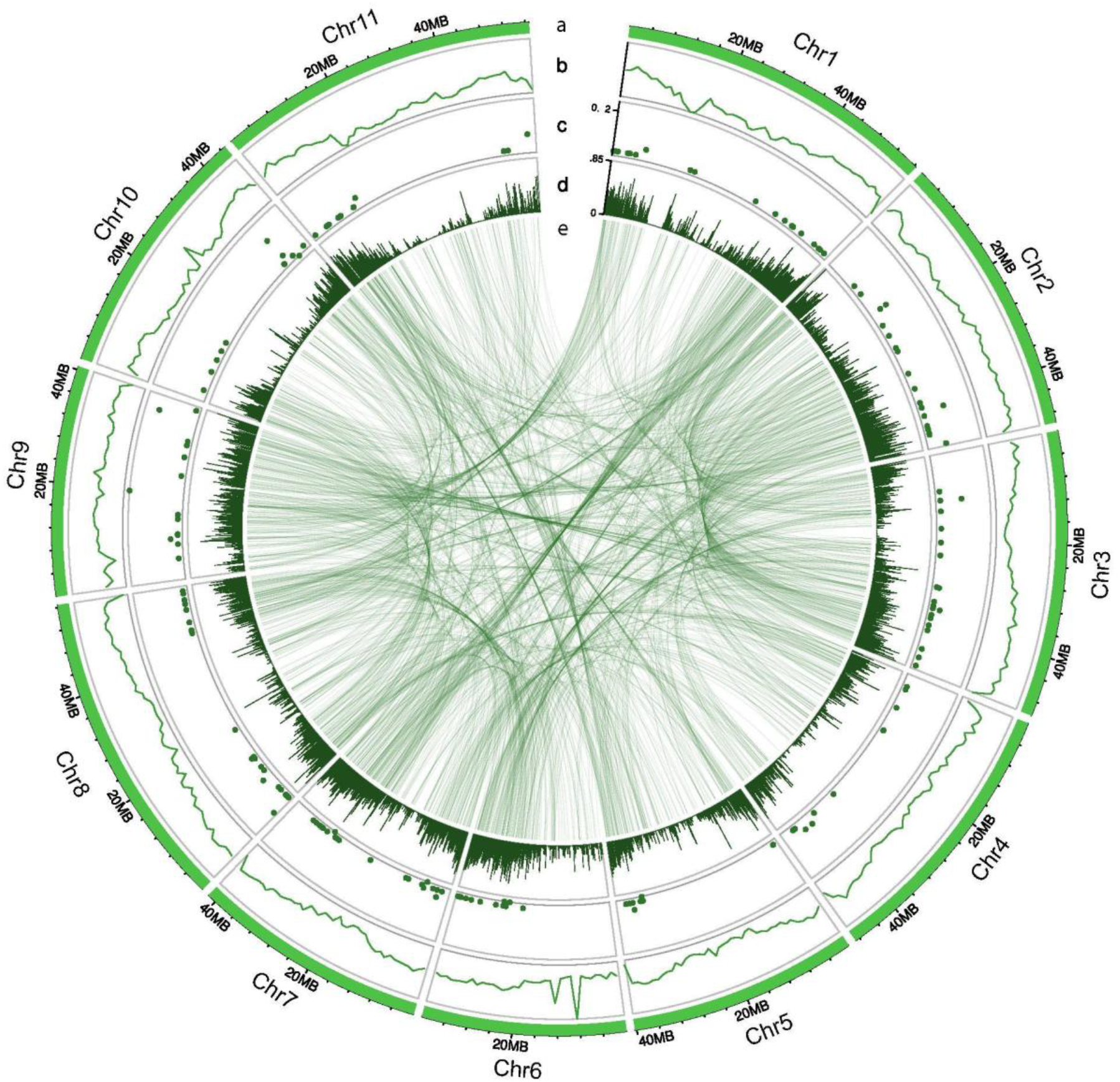
Circos plot illustrating the genomic landscape and structural features of the *P. vulgaris* Negro Jamapa chromosomes. Track (a) displays chromosomal ideograms, arranged sequentially and scaled according to chromosome length. Track (b) represents the distribution of repetitive elements (green lines). Track (c) shows the genomic positions of micro-RNA (miRNA) precursor genes (dark green points). Track (d) corresponds to annotated protein-coding genes density (vertical green lines). (e) Colored inter-chromosomal links connect pairs of potentially duplicated genes, representing putative segmental duplications or paralogous relationships.

Overall, these results offer a comprehensive view of the Negro Jamapa genome and illustrate how repetitive sequences, gene organization, and large-scale duplications have shaped the evolution of this *P. vulgaris* cultivar.

### Structural Variation Between Negro Jamapa and G19833

To contextualize the quality of the Negro Jamapa assembly and identify cultivar-specific structural features relevant for downstream functional genomics, we performed a comparative analysis against the widely used G19833 reference genome. Whole-genome alignment revealed high levels of structural and sequence variation, showing the deep divergence between the two main gene pools of common beans **(Figure 2A)**. In the aligned regions, the SNP rate was estimated at 7.6 per kilobase, which agrees with previous reports about strong nucleotide differentiation between Mesoamerican and Andean lineages. However, structural variants represented the majority of the genomic differences, with several categories detected **(Figure 2B)**. Among them, highly diverged regions represented the largest proportion (23.4%) of the polymorphic DNA, followed by unaligned regions in the query genome (23.2%) and inter-chromosomal translocations (20.6%). Inversions accounted for 12.7%, while copy number variants, including copy gains (6.2%) and segmental duplications (5.4%), represented together more than 11%. Smaller fractions were related to insertions (5.2%) and tandem repeats (0.6%).

**Figure 2.**
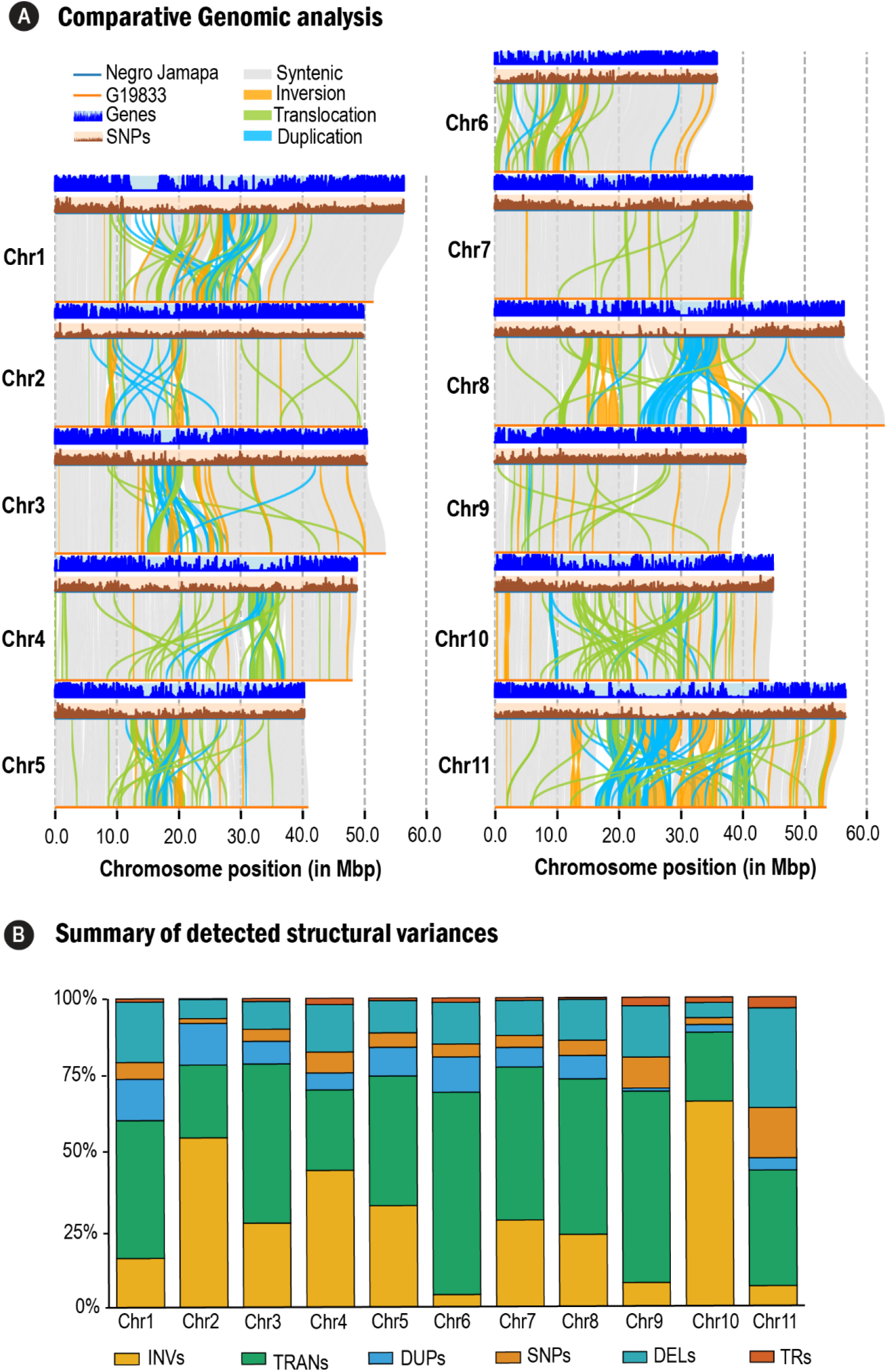
Genome-wide distribution of structural variants between *P. vulgaris* Negro Jamapa and G19833 genomes. A. Comparative genomic analysis. For each chromosome, the upper track displays the distribution of SNPs (blue), and the lower track represents annotated genes (brown). Vertical links between chromosomes indicate structural variants identified by SyRI and visualized with plotsr, including inversions (yellow), translocations (green), and duplications (light blue). Syntenic regions are represented in grey. Vertical dashed lines indicate the relative chromosome position (Megabases). B. Summary of detected structural variants. Chromosomal representation of the cumulated length proportion of different classes of structural variants identified across the 11 chromosomes of P. *vulgaris* Negro Jamapa. Each bar indicates the relative abundance (0–100%) of variant types per chromosome. INVs: inversions; TRANs: translocations; DUPs: duplications; SNPs: single nucleotide polymorphisms; DELs: deletions; TRs: tandem repeats. Structural variants were detected using SyRI (Synteny and Rearrangement Identifier) based on pairwise genome alignment.

In general, these results indicate that large structural changes and highly diverged genomic regions are the main components of divergence between Negro Jamapa and G19833, more than single-nucleotide changes. Such extensive structural differences probably contribute to reproductive isolation, local adaptation, and the phenotypic differences between these two main *P. vulgaris* gene pools.

### Phabase: an integrative database for Phaseolus spp

*Phabase* was implemented as a modular web system designed to integrate genomic and transcriptomic information from *P. vulgaris*. The architecture follows a three-tier model comprising the database layer, web services layer, and presentation layer, each deployed in isolated Docker containers to ensure scalability and maintainability **(Figure 3)**. Overall, *Phabase* brings together genomic and transcriptomic tools within a streamlined, user-friendly workflow **(Figure 4)**. The platform provides open access to multiple datasets, including *P. vulgaris* G19833 v2.1 (Schmutz *et al*., 2014), *P. vulgaris* BAT93 v1.0 (Vlasova *et al*., 2016), and the newly assembled *P. vulgaris* cv. Negro Jamapa v1 generated in this study (**Figure 4A**). It is an intuitive platform for exploring genes and retrieving both nucleotide and amino acid sequences from any of the available genomes **(Figure 4B)**. Additionally, *Phabase* integrates the Basic Local Alignment Search Tool (BLAST), enabling users to identify orthologous sequences across the available genomes using either nucleotide or amino acid queries **(Figure 4C)**.

**Figure 3:**
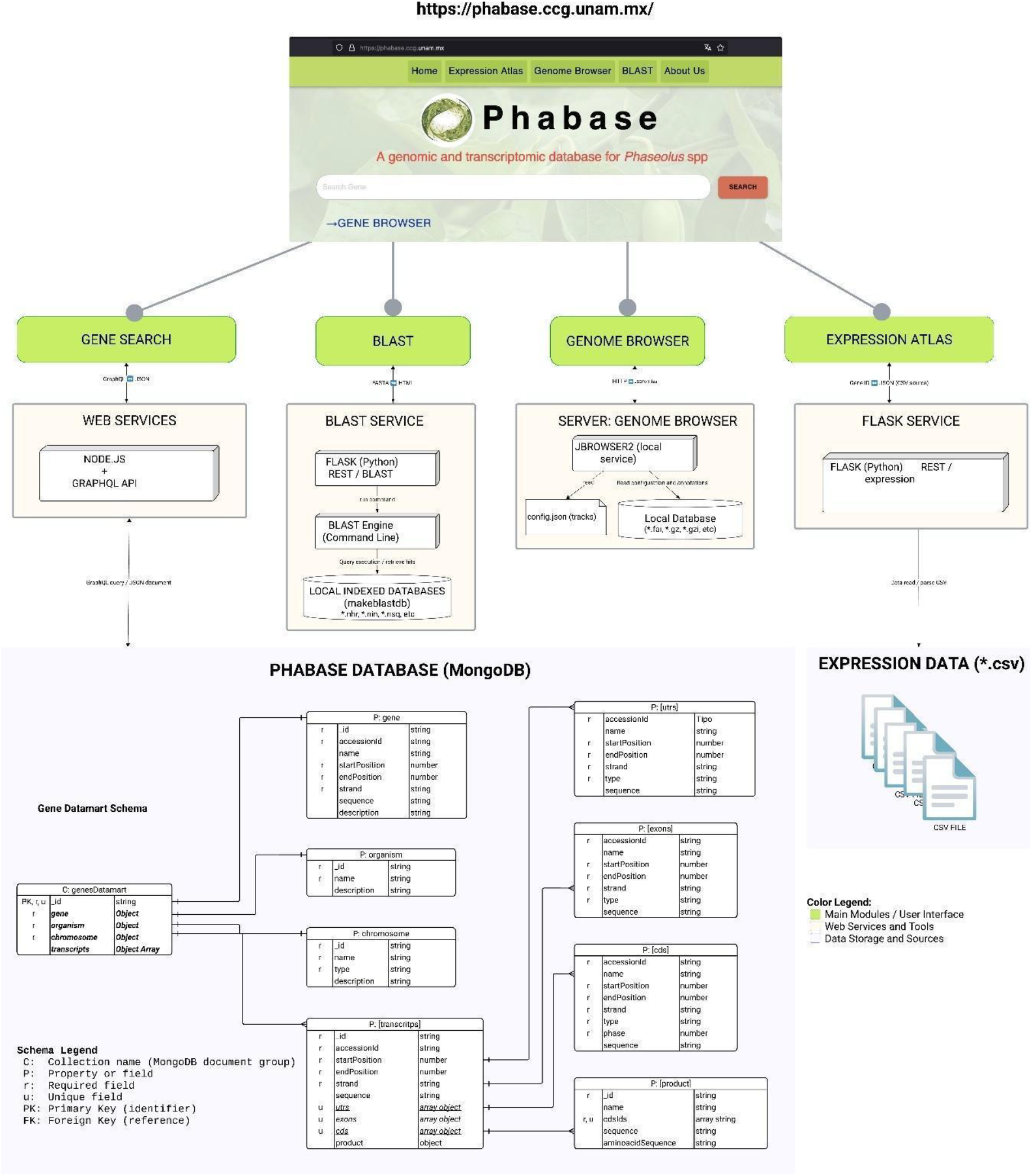
System architecture of *Phabase*. The diagram illustrates the modular architecture of the *Phabase* platform, showing the interaction between the main user modules, web services, and data sources. Green boxes represent the user interface modules, orange boxes correspond to web services and computational tools, and blue boxes denote data storage and sources.

**Figure 4.**
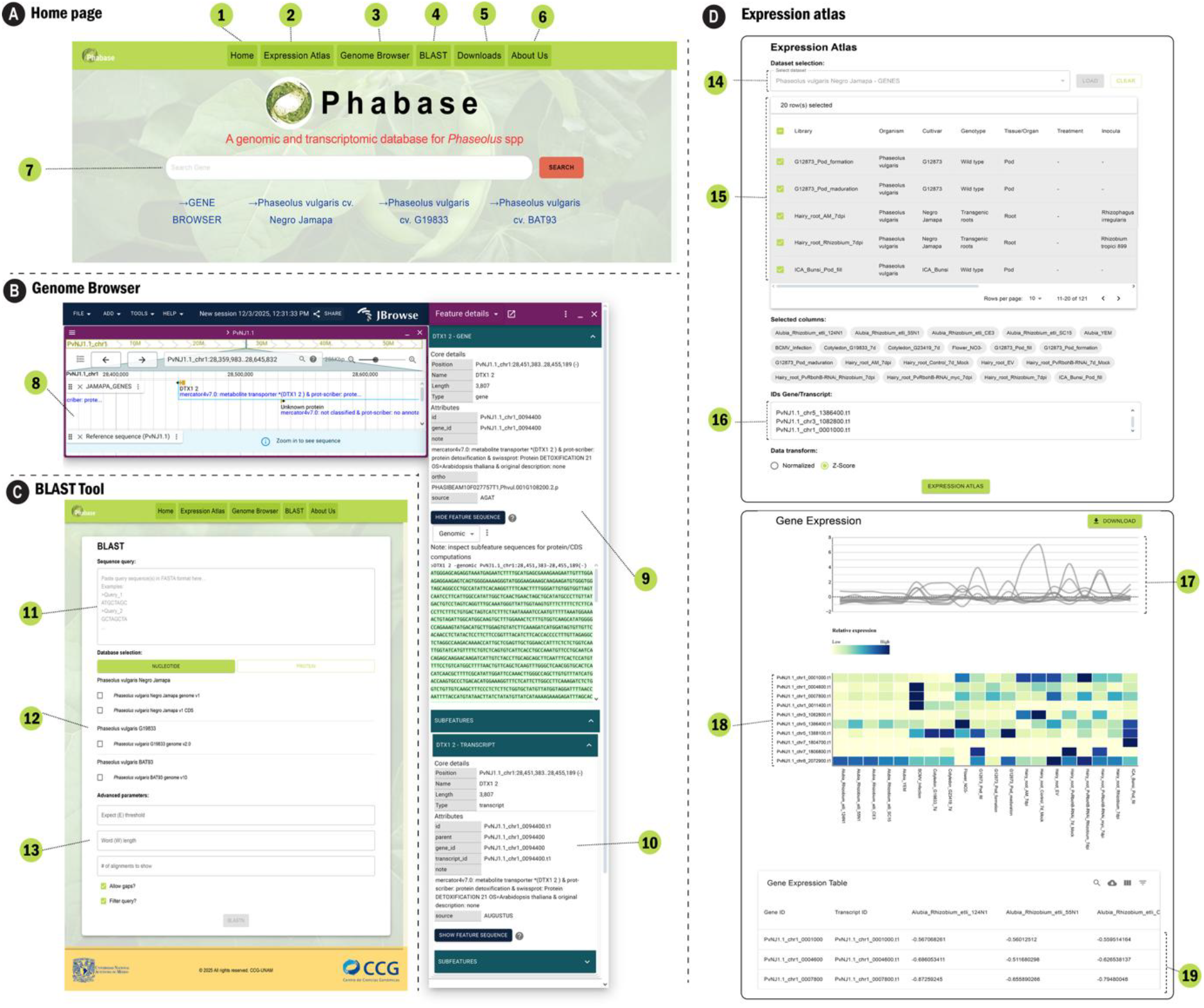
Overview of *Phabase* tools and interface. **(A)** Home page of *Phabase* showing the main navigation menu. *Home* (1), *Gene Expression* (2), *Genome Browser* (3), *BLAST* (4), and *About Us* (5) sections. The search bar (6) allows users to query genes of interest directly. **(B)** Genome Browser interface illustrates the visualization of genomic regions. Genes are represented as annotated tracks and users can inspect gene models, annotations, and neighboring genomic elements (7). Selecting a feature opens a detailed information panel (8), which includes gene coordinates, structure, functional annotation, and nucleotide sequences. Additional transcript-level annotations and predicted subfeatures (9) can be explored through collapsible sections. **(C)** BLAST interface allows users to submit a nucleotide or protein query sequence in FASTA format (10) and search against multiple Phaseolus reference genomes selectable from the database list (11). Optional advanced parameters (12), including E-value threshold, word size, number of alignments to display, and filtering options, enable customized similarity searches.

### Expression atlas for genes and microRNAs

In order to improve community access to transcriptomic data analysis and have a wider panorama of gene and microRNA expression in *P. vulgaris*, we collected transcriptomic and micro-transcriptomic data from several studies and organized them into an integrated expression atlas. For the gene expression part, the atlas now includes 121 experimental conditions that cover a wide range of biological conditions **(Figure 5A)**. Around 52% of them are related to plant development, while about 48% correspond to stress conditions, such as drought, pathogen infection, abiotic stress, and symbiotic interactions **(Figure 5A)**. The datasets come from 21 cultivars, with Negro Jamapa representing more than 36% of all samples **(Figure 5A)**. The data includes 10 different organs or tissues, where roots are the most represented (52% of samples) **(Figure 5A)**. These studies were led by research groups from nine countries, mainly from the United States of America (36%) and Mexico (21%), which shows the strong international interest in common bean research, especially in relation to stress and development **(Figure 5C)**.

**Figure 5.**
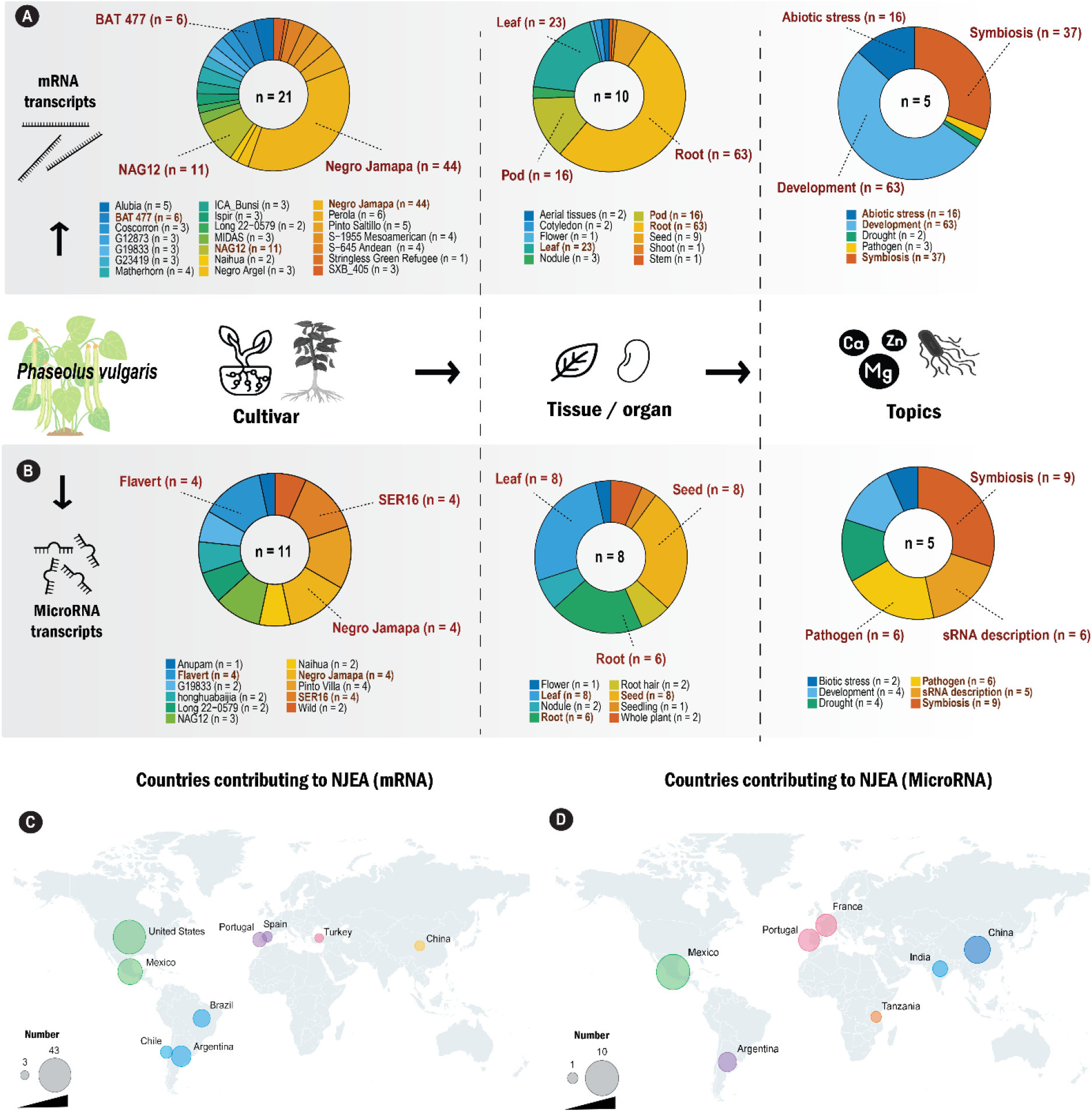
Overview of the Expression Atlas module in *Phabase*. Summary of the datasets integrated into *Phabase* for **(A)** Gene and **(B)** microRNA Expression interface. Charts display the number of samples categorized by cultivar, tissue/organ, and topic. The number at the center of each chart indicates the total number of elements represented in the corresponding chart. Geographic distribution of the research groups that generate datasets that were processed to obtain the **(C)** Gene and **(D)** microRNA Expression module. The diameter of a circle is proportional to the number of conditions produced in the corresponding country. Colors differentiate regions of the world

For the microRNA expression atlas, 284 small RNAs have been selected. Almost all of them are microRNAs (274), including 154 non-common bean-specifics and 120 only identified in *P. vulgaris* genome (Formey *et al*., 2015; Formey *et al*., 2016). The other ten small RNAs are tRFs, transfer RNA (tRNA)-derived small RNA fragments from rhizobia (Ren *et al*., 2019). In this atlas, we gather data from 30 experimental conditions. The most common topics were focused on plant-microbe interactions, particularly on symbiotic interactions (30%) and pathogen responses (20%) **(Figure 5B)**. In this case, the datasets come from 11 cultivars, including Negro Jamapa, Flavert, and Pinto Villa (each one representing around 13% of all samples) **(Figure 5B)**. Expression profiles were obtained from eight organs or tissues, mainly leaves (27%), seeds (27%), and roots (20%) **(Figure 5B)**. These studies were produced by projects led by research groups from seven countries, with an important contribution from Mexico (33%), reflecting a leading role in small RNA studies in beans **(Figure 5D)**.

Altogether, this atlas represents the most complete and diverse collection of *P. vulgaris* expression data so far. It offers a strong base for comparative studies, helps to identify regulatory networks, and facilitates the analysis of gene/microRNA interactions across different cultivars, organs, and environmental conditions.

### Exploring MYB36 orthologs in P. vulgaris Negro Jamapa using Phabase

To illustrate how *Phabase* can be applied by typical end users to explore gene function and generate data-driven hypotheses, we selected MYB36 as a case study (**Figure 6A)**. This is a transcription factor recently characterized for its role in Casparian strip formation and its indirect influence on microbial colonization dynamics in *Arabidopsis thaliana* (Tsai *et al*., 2025).

**Figure 6.**
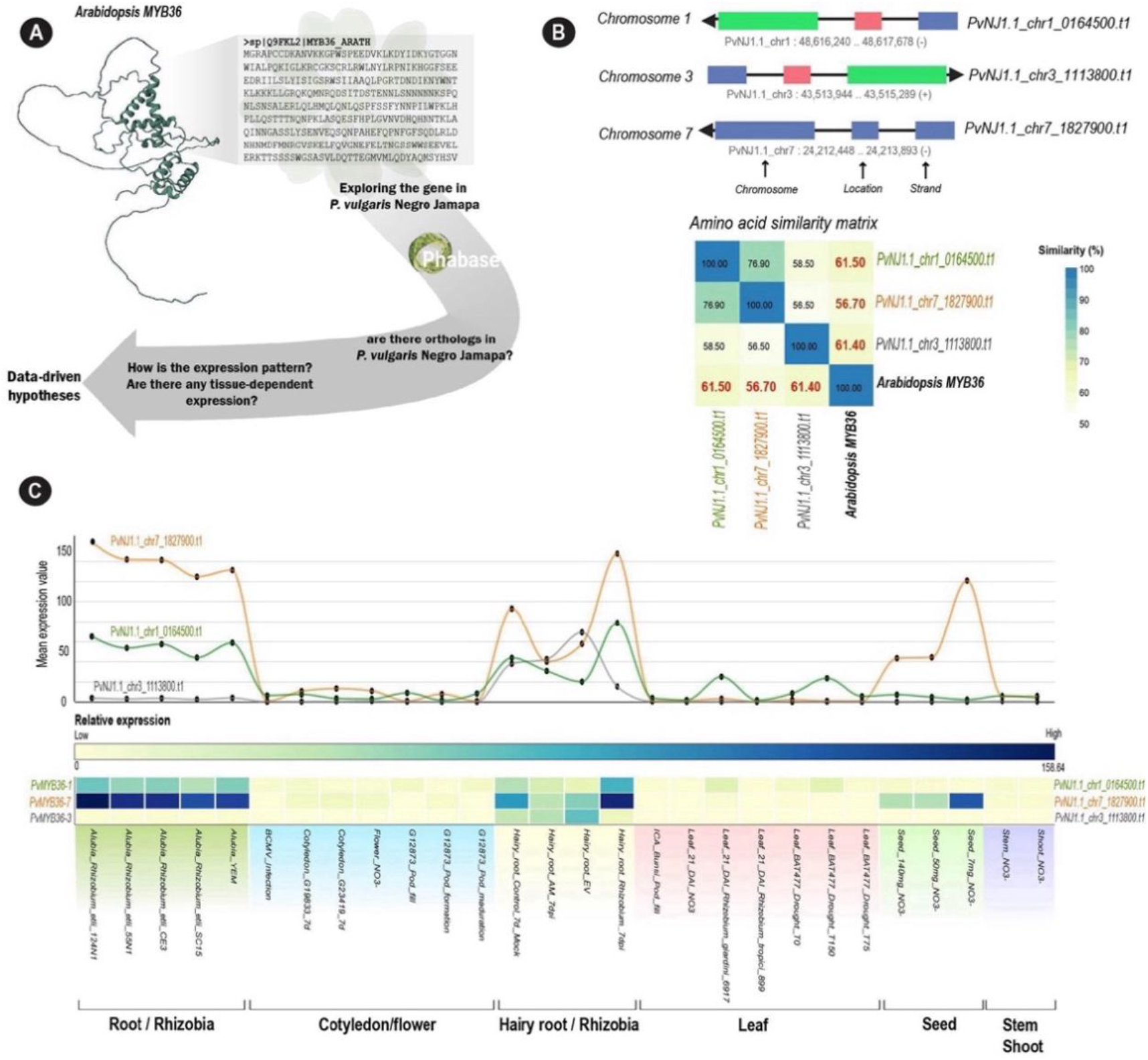
*Phabase*-enabled identification of MYB36 orthologs in *P. vulgaris* cv. Negro Jamapa. **(A)** Workflow illustrating how *Phabase* was used to identify MYB36 orthologs in *P. vulgaris* cv. Negro Jamapa. **(B)** Genomic locations and amino acid similarity matrix of the putative MYB36 orthologs. **(C)** Expression profiles of the identified MYB36 orthologs across different tissues and conditions.

We first investigated whether orthologous sequences of *AtMYB36* are present in the *P. vulgaris* cv. Negro Jamapa genome. To do so, we retrieved the *A. thaliana* MYB36 amino acid sequence from the UniProt database and used it as a query in the *Phabase* BLASTP tool against the *P. vulgaris* Negro Jamapa v1 protein dataset. This search identified three candidate sequences with E-values below 4e-31: PvNJ1.1_chr7_1827900.t1 (Identities = 55/60, 92%; Positives = 59/60, 98%; Gaps = 0%), PvNJ1.1_chr3_1113800.t1 (Identities = 54/60, 90%; Positives = 57/60, 95%; Gaps = 0%), and PvNJ1.1_chr1_0164500.t1 (Identities = 55/60, 92%; Positives = 59/60, 98%; Gaps = 0%), located on chromosomes 7, 3, and 1, respectively (**Figure 6B**). Overall, these three putative orthologs shared more than 56% sequence identity with *A. thaliana* MYB36 (**Figure 6B)** and were annotated as MYB transcription factors according to Mercator v4 pipeline (Bolger *et al*., 2021).

In *A. thaliana, MYB36* (AT5G57620) is primarily expressed in roots and seeds during germination (Klepikova *et al*., 2016; Reiser and et, 2022). We hypothesized that the identified orthologs in *P. vulgaris* would exhibit a similar expression pattern. To investigate this, we analyzed the expression profiles of the putative MYB36 orthologs (*PvNJ1*.*1_chr7_1827900*.*t1* = *Pv*MYB36-7, *PvNJ1*.*1_chr3_1113800*.*t1* = *Pv*MYB36-3, and *PvNJ1*.*1_chr1_0164500*.*t1* = *Pv*MYB36-1) across various tissues and experimental conditions using the NJEA module in *Phabase* (**Figure 6C)**. Overall, the three orthologs were predominantly expressed in roots, with *PvMYB36-7* showing the highest transcript abundance. Notably, *PvMYB36-7* was also the only ortholog detected in seeds. Its expression profile, characterized by strong root-specific expression and moderate activity in developing seeds (**Figure 6C)**, closely mirrors that of *MYB36* in *A. thaliana*, which is known to regulate endodermal differentiation and Casparian strip formation. This similarity in spatial expression patterns suggests that *Pv*MYB36-7 likely represents the functional MYB36 ortholog in *P. vulgaris*, potentially playing a conserved role in root barrier formation and developmental regulation.

Together, this quick analysis illustrates the utility of *Phabase* for integrating sequence-based ortholog identification with expression profiling to rapidly generate data-driven hypotheses.

## Discussion

Negro Jamapa represents a key cultivar for Mesoamerican common bean breeding and genomic research. It is not only part of the national food security pilar of Mexico but also represents a key model for functional genomics analyses and the generation of expression data. The assembly and annotation of its genome, one of the aims of this project, give a strong base to study the genomic structure and functional diversity of Mesoamerican common beans. This new genome helps to close a gap in bean genomics, since accessible genomic analysis tools for the common bean community are lacking and only available for the G19833 genome reference, from the Andean gene pool. With this high-quality assembly and the creation of a unified data portal, *Phabase*, our work improves the access and integration of genomic, transcriptomic, and small RNA data, from an agronomically important Mesoamerican cultivar, for the bean research community.

### Comparative Genome Architecture and Evolutionary Insights

The Negro Jamapa genome shows robust quality and completeness (N50 = 45 Mb; 98.4%), which are higher than those of the reference genome G19833 (N50 = 2 Mb; 97.7%). This difference reflects how long-read sequencing and assembly technologies allow the generation and democratization of high-quality genomes, especially for complex plant genomes. Other studies show that structural variations contribute to reproductive isolation, local adaptation, and domestication traits in *P. vulgaris* (Ambachew *et al*., 2024; MacQueen *et al*., 2022; Rendon-Anaya *et al*., 2017; Schmutz *et al*., 2014). As expected, Negro Jamapa and G19833 belonging to different gene pools, their genomes present clear divergence, mainly explained by large structural variants. The facilitating access to this information allows the community to explore the genomic details of this divergence and reveal the phenotypic and physiological characteristics specific to Negro Jamapa and Mesoamerican common beans.

### An Integrative Atlas for Transcriptomic and Small RNA Data

To extend the utility of the Negro Jamapa genome for functional and comparative studies, we developed *Phabase*, an open and integrative data portal designed for the *P*. research community. *Phabase* offers a complete expression atlas that integrates both coding and non-coding genes, connecting genomic information with transcriptomic and small RNA data.

The expression section currently includes 121 experimental conditions representing diverse biological contexts, such as plant development, biotic and abiotic stresses, and symbiotic interactions, across 21 cultivars, increasing five times the quantity of information compared to the original gene atlas (O’Rourke *et al*., 2014). On average, these experiments have also more than two replicates, increasing the robustness of the data, compared to the one-replicate experiments available originally (O’Rourke *et al*., 2014). Root samples dominate the collection (around 51%), highlighting the strong focus of the bean community on root biology, nodulation, and Rhizobial symbiosis. Remarkably, Negro Jamapa contributes to more than one-third of the total data, reinforcing the importance of this cultivar in the community and the need to generate robust genomic data for this cultivar.

For small RNAs, *Phabase* compiles data of 284 small RNAs, from 30 experimental conditions involving 11 cultivars, with samples from different organs and stress environments. To our knowledge, this is the first publicly available plant sRNA expression atlas website and the unique to include exogenous sRNA identified as host regulators by cross-kingdom RNAi (Formey *et al*., 2016). Together, these efforts make *Phabase* the most comprehensive and diverse transcriptomic resource currently available for *Phaseolus*.

*Phabase* is not only as a data repository but also a platform for functional exploration. We included tools for visualization of expression profiling and comparative analyses, allowing users to easily explore the data and test hypotheses without bioinformatics skills. *Phabase* also helps to reduce bean genomic data fragmentation by integrating and homogenizing the publicly available data that were previously dispersed across different sources. We believe that it creates a unified and accessible space for large-scale comparative studies and meta-analyses in bean genomics.

As proof of concept and illustrate how *Phabase* can help when a user is interested in the analysis of one specific protein from another organism, we looked at AtMYB36, a transcription factor known for its role in the Casparian strip and endodermis formation in *Arabidopsis thaliana*. When we searched for similar sequences in the Negro Jamapa genome, we found three possible candidates: *PvMYB36-7, PvMYB36-3, and PvMYB36-1*, each one on a different chromosome. Investigating its expression profile, we confirmed that *PvMYB36-7* could be the functional orthologs of *AtMYB36*. These two genes display similar expressions in roots, with some signal in developing seeds. This small test shows that, thanks to *Phabase*, the community can quickly check homology, look at expression across several conditions and cultivars, and start forming hypotheses without bioinformatic skills.

The Negro Jamapa genome and *Phabase* together fill a gap for the *Phaseolus* community: a Mesoamerican reference genome with high contiguity and a platform gathering different types of genomic data. Compare Mesoamerican and Andean gene pools and explore questions related to domestication, stress responses, or plant-microbe interaction is now more accessible. More generally, these tools reduce the fragmentation of bean genomic data and make it easier for researchers, especially those without deep bioinformatics experience, to work with large-scale datasets.

## Materials and methods

### Biological material and sequencing

Commercial seeds of *P. vulgaris* var. Negro Jamapa were disinfected with 3% sodium hypochlorite for 10 minutes, followed by three washes with sterile distilled water to eliminate residual chlorine. The sterilized seeds were sown in pots containing Peat Moss and Leica substrate (ratio 3:1) and cultivated under greenhouse conditions for two weeks. Young leaves from 15 individual plants were harvested, frozen, and subsequently macerated in liquid nitrogen. High molecular weight DNA was extracted using the NucleoBond HMW DNA kit (Macherey Nagel), in accordance with the manufacturer’s protocol. The resulting HMW DNA was submitted to Novogene for sequencing via Hi-Fi PacBio technology, yielding 95.3 Gb sequencing data with the following parameters: Maximum read length, 64 kb; Mean read length, 21 kb; N50 read length, 21 kb.

### Assembly pipeline

HiFi PacBio reads were assembled to contigs using hifiasm version 0.20.0-r639 with the default parameters (Cheng *et al*., 2021). Organellar genomes were assembled using Oatk version 1.0 (minimum k-mer coverage = 200 and k-mer length = 1001) (Zhou *et al*., 2025). We then performed homology searches for hifiasm contigs over the organellar genomes using BLAST. Short hifiasm contigs (< 1,000,000 nt) with high similarity (BLAST search filters: e-value < 1E-06, % identity > 99%, match length > 1000) to the organellar genomes were removed, leaving the contigs pertaining to the nuclear genome. These nuclear contigs then were subjected to the correcting and scaffolding steps of the RagTag pipeline (Alonge *et al*., 2022). Genome assembly *P. vulgaris* G19833 v2.0 was implemented in these steps as the reference genome (Schmutz *et al*., 2014). The final assembly, including all 11 chromosomes that were placed to the reference genome, the unplaced contigs, and the organellar genomes were then subjected to gene prediction.

### Genome annotation

The repeat sequences in the genome were identified using Repeatmodeler (v2.0.5) (Flynn *et al*., 2020) and were masked using RepeatMasker (v4.1.5) (Tarailo-Graovac and Chen, 2009). Previously published RNAseq (O’Rourke *et al*., 2014) data were mapped using STAR (v2.7.11b) (Dobin *et al*., 2013). The Viridiplantae OrthoDB (v10.1) (Kriventseva *et al*., 2019), merged with the proteins from the reference genome of *P. vulgaris* were also used in the pipeline. The annotation was done using Braker3 (v3.0.8) (Gabriel *et al*., 2024). The completeness of the annotations was assessed using BUSCO (v5.8.3) (Manni *et al*., 2021). The protein sequences were functionally annotated using Mercator4 v7.0 (Bolger *et al*., 2021). The UTRs for the genes were added using Stringtie based on the RNAseq data mapping (Shumate *et al*., 2022). miRNA genes were predicted thanks to miRDeep-P2 (Kuang *et al*., 2019) and Shortstack (Axtell, 2013); and validated by miRScore (Vanek *et al*., 2025), using the dataset displays in the miRNA Gene Atlas as sRNA library inputs (Table S1).

### Overview and technologies Phabase

The database layer is based on MongoDB, which stores the integrated data model (genesDatamart) derived from multiple collections containing gene, transcript, and product annotations. This hierarchical structure enables fast retrieval of gene-centric information including sequence data, genomic coordinates, and functional annotations. The web services layer is implemented in Node.js and exposes a GraphQL API for handling data queries and gene search operations. This layer also communicates with specialized Flask (Python) services responsible for processing computational requests. The presentation layer is built with Next.js, providing an interactive user interface named JAMAPA Browser. This interface integrates four functional modules: (i) Gene Search, (ii) Expression Atlas, (iii) Genome Browser, and (iv) BLAST. All client requests are routed through a Nginx reverse proxy, which manages HTTPS communication between the user and the web services. The modular design allows each component to operate independently, facilitating system updates and integration of new datasets or analysis tools. All components communicate through REST or GraphQL interfaces, ensuring reproducibility, interoperability, and transparent data access for both visualization and downstream analysis.

The Genome Browser module uses JBrowse 2, deployed as a separate service configured via a JSON file with references to genome FASTA and GFF3 annotation files. In addition, *Phabase* integrates the Basic Local Alignment Search Tool (BLAST), enabling users to identify orthologous sequences across the available genomes using either nucleotide or amino acid queries. The BLAST service runs locally on a dedicated server, where the Flask API executes the BLAST command-line tool against locally indexed databases generated with makeblastdb. Search results are returned in JSON or tabular format. For more precise control, advanced parameters are enabled, allowing users to adjust their searches according to specific needs.

### Expression Atlas

The Expression Atlas service accesses precomputed RNA-seq data stored in CSV files. Flask reads and parses these datasets, serving the results as JSON objects that are visualized dynamically in the front-end using D3.js. The gene and microRNA expression atlas encompass 261 libraries from 121 experimental conditions, and 67 libraries from 30 experimental conditions (Hiz *et al*., 2014, O’Rourke *et al*., 2014, Wu *et al*., 2014, Dalla Via *et al*., 2015, Formey *et al*., 2015, Formey *et al*., 2016, Martin *et al*., 2016, Nanjareddy *et al*., 2017, Sosa-Valencia *et al*., 2017, Wu *et al*., 2017, Singh *et al*., 2018, Fonseca-Garcia *et al*., 2019, Gregorio Jorge *et al*., 2020, Pereira *et al*., 2020, Leitao *et al*., 2021, Mwaipopo *et al*., 2021, Parreira *et al*., 2021, Castaingts *et al*., 2022, Clua *et al*., 2022, Parker *et al*., 2022, Sanchez-Correa *et al*., 2022, Lopez *et al*., 2023, Darrasse *et al*., 2024, Tapia *et al*., 2024, Wu *et al*., 2024), respectively, including responses to abiotic stress, symbiotic interactions, and various developmental stages (Table S1 and S2) NJEA allows users to visualize and compare gene expression patterns across diverse conditions, facilitating the identification of context- and tissue-specific gene regulation. To build the NJEA, raw RNA-seq reads were retrieved from NCBI using SRA-toolkit (Leinonen *et al*., 2011) and low-quality bases and adapter sequences were removed with fastp (Chen *et al*., 2018). The resulting high-quality reads were aligned to the *P. vulgaris* Negro Jamapa genome using the splice-aware aligner STAR (Dobin *et al*., 2013). Transcript-level expression values were quantified with featureCounts (Liao *et al*., 2014) and read counts were subsequently normalized using edgeR (Robinson *et al*., 2010). For miRNA, reads where collapsed and counted using FASTQ/A Collapser from the FASTX-Toolkit (http://hannonlab.cshl.edu/fastx_toolkit/) and annotated with compiled data from previous studies(Formey *et al*., 2015; Formey *et al*., 2016; Ren *et al*., 2019). Given that the integrated datasets span multiple cultivars, growth facilities, sampling time points, and distinct biological questions, global batch-effect correction was not applied, as it would risk confounding technical variation with genuine biological signal. Phabase is therefore best understood as an exploratory resource for the community — a tool to survey expression tendencies, generate hypotheses, and identify candidates of interest across diverse conditions — rather than a platform for precise quantitative expression measurements. Users are strongly encouraged to carefully select contextually comparable experiments when performing cross-experiment analyses.

## Data availability

Phabase and corresponding data can be accessed at the following URL: https://phabase.ccg.unam.mx/download and Bioproject: PRJNA1428309.

## Code availability

All the scripts used in this study are deposited on GitHub: https://github.com/tyakyol/Phabase and https://github.com/phabasedb

## Acknowledgements

The authors thank the Bioinformatics Analysis Unit (UAB) and the Technology and Information Management Unit (UATI) for their invaluable assistance with genome assembly and website management.

